# A direct proof that sole actin dynamics drive membrane deformations

**DOI:** 10.1101/307173

**Authors:** Camille Simon, Rémy Kusters, Valentina Caorsi, Antoine Allard, Majdouline Abou-Ghali, John Manzi, Aurélie Di Cicco, Daniel Lévy, Martin Lenz, Jean-François Joanny, Clément Campillo, Julie Plastino, Pierre Sens, Cécile Sykes

## Abstract

Cell membrane deformations are crucial for proper cell function. Specialized protein assemblies initiate inward or outward membrane deformations that turn into, for example, filopodia or endocytic intermediates. Actin dynamics and actin-binding proteins are involved in this process, although their detailed role remains controversial. We show here that a dynamic, branched actin network is sufficient, in absence of any membrane-associated proteins, to initiate both inward and outward membrane deformation. With actin polymerization triggered at the membrane of liposomes, we produce inward filopodia-like structures at low tension, while outward endocytosis-like structures are robustly generated regardless of tension. Our results are reminiscent of endocytosis in mammalian cells, where actin polymerization forces are required when membrane tension is increased, and in yeast, where they are always required to overcome the opposing turgor pressure. By combining experimental observations with physical modeling, we propose a mechanism for actin-driven endocytosis under high tension or high pressure conditions.

---

Many cell functions rely on the ability of cells to change their shape. The deformation of the cell membrane is produced by the activity of various proteins that curve the membrane inwards or outwards, by exerting pulling and pushing forces or by imposing membrane curvature via structural effects. Membrane invagination (or inward deformation of the cell membrane) can be initiated by specific proteins, such as clathrin, which coat the membrane and impose geometrical constraints that bend the membrane inwards. In this view, the action of the actin cytoskeleton, a filamentous network that forms at the membrane, is crucial only at a later stage for membrane elongation. Nevertheless, impressive correlation methods revealed unambiguously that, in yeast, membrane bending is not triggered by the presence of coat proteins, but by a dynamic actin network formed at the membrane through the Arp2/3 complex branching agent ^1–3^. In mammalian cells, clathrin-mediated endocytosis requires the involvement of actin if the plasma membrane is tense, *e.g*. following osmotic swelling or mechanical stretching ^4^. However, the exact mechanism of membrane deformation in this process is still poorly understood. Strikingly, the same type of branched actin network is able to bend the membrane the other way in dendritic filopodia, outward-pointing membrane deformations that precede the formation of dendritic spine in neurons ^5^. Dendritic filopodia appear different from conventional filopodia where actin filaments are visibly parallel. The ability of a branched actin network to produce a filopodia-like membrane deformation has never been investigated.

How the same branched actin structure can be responsible for the initiation of filopodia, which are outward-pointing membrane deformations, as well as endocytic cups that deform the membrane inward, is what we want to address in this paper. Such a question is difficult to investigate in cells that contain redundant mechanisms for cell deformation. Actin dynamics triggered at a liposome membrane provide a control on experimental parameters such as membrane composition, curvature and tension, and allow the specific role of actin dynamics to be addressed. We evidence that the same branched actin network is able to produce both endocytosis-like and filopodia-like deformations. With a theoretical model, we predict under which conditions the stress exerted on the membrane might lead to inward and/or outward pointing membrane deformations. Combining experiments and theory allows us to decipher how the interplay between membrane tension and actin dynamics produces inward or outward membrane deformations.

## Membrane deformations:tubes and spikes

Liposomes covered with an activator of the Arp2/3 complex, SpVCA, are placed in a mixture containing monomeric actin, profilin, the Arp2/3 complex and capping protein (CP) in order to grow a branched actin network at their surface (Materials and Methods and Fig. 1A). Strikingly, imaging the membrane of liposomes in the presence of a growing actin network reveals that the liposome surface is not smooth, but instead shows a rugged profile. Indeed, membrane tubes, hereafter called “tubes”, are observed to radiate from the liposome surface and extend into the actin network (Fig. 1B), even when comet formation has occurred ^6,7^ (Supplementary Fig. 1). Interestingly, some liposomes display another type of membrane deformation, characterized by a conical shape that points towards the liposome interior, hereafter referred to as “spikes” (Fig. 1B). Some of the liposomes carry both tubes and spikes, while others display neither, despite the presence of an actin network at the membrane (Fig. 1B).

**Figure 1:**
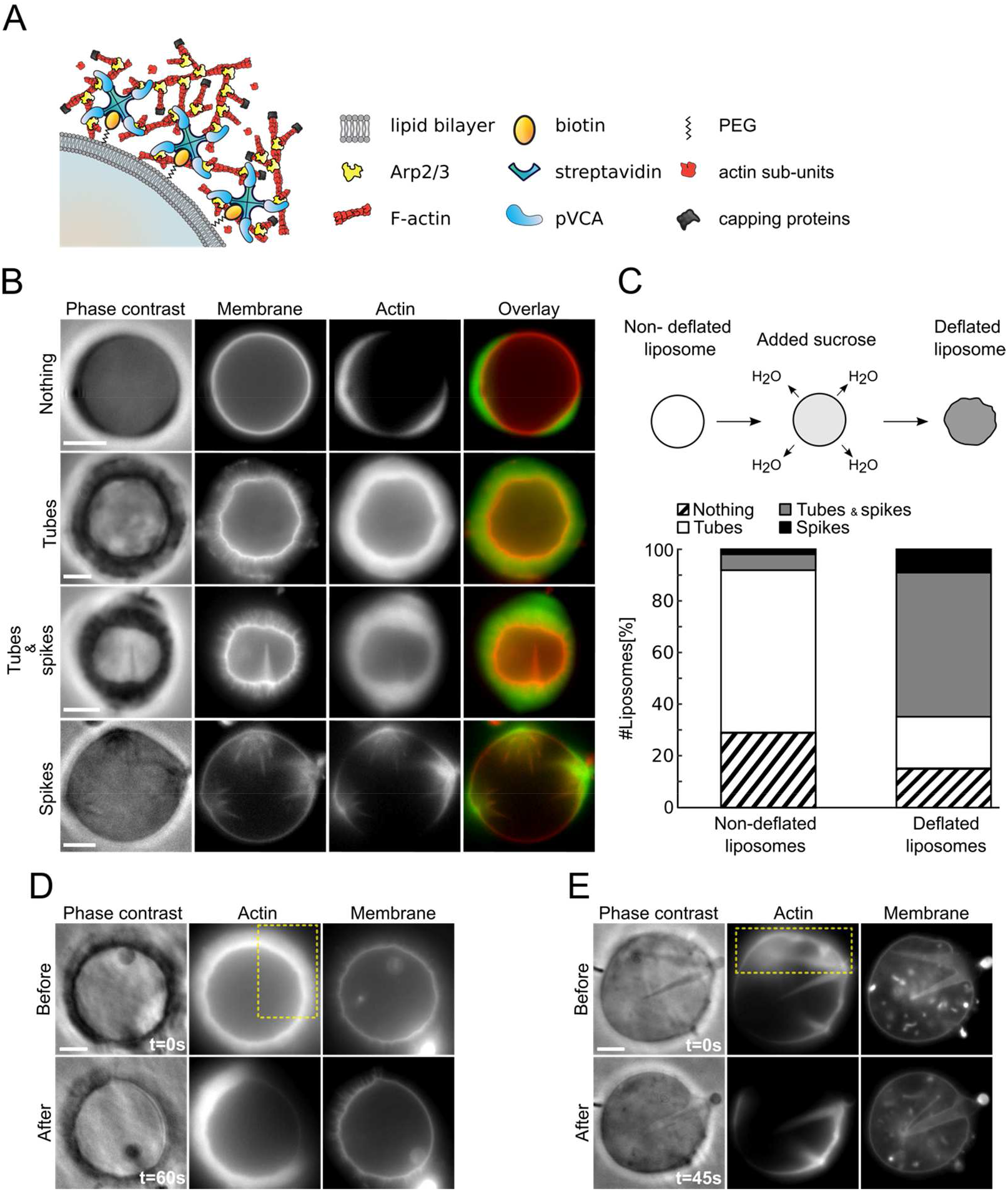
Experimental system and observations. (A) Scheme of the experimental system; proteins not to scale. (B) Membrane deformations in both non-deflated (three first rows) and deflated conditions (last row). (C) Top: liposome deflation. Bottom: number of liposomes displaying nothing, tubes, both tubes and spikes, and only spikes. Non-deflated liposomes, n=311. Deflated liposomes, n=123. (D, E) Actin network photo-damage (yellow dashed rectangle) on a liposome displaying membrane tubes (D) or spikes (E). Phase contrast and epifluorescence microscopy of membrane and actin network. Scale bars, 5μm.

We now address the role of membrane tension on the appearance of tubes and spikes. Under conditions of normal osmotic pressure (200 mOsm), 63.0% of liposomes display tubes only, 2.3% show spikes only, while 6.1% of liposomes have a mix of both, and 28.6% have neither (Fig. 1C). To examine how membrane tension affects the occurrence of tubes and spikes, liposomes are deflated by increasing the osmotic pressure of the working buffer to 400 mOsm. On the one hand, a huge increase in the number of liposomes displaying spikes is observed when membrane tension is lowered in deflated liposomes. Indeed 65.0% of deflated liposomes display spikes (with or without tubes), compared to 8.4% in non-deflated conditions (Fig. 1C, p < 0.0001). On the other hand, the frequency with which tubes are observed is essentially unaffected by a change in membrane tension: 69.1% for non-deflated liposomes compared to 74.8% for deflated liposomes (not significant, p = 0.24 > 0.05). The presence of membrane tubes and spikes clearly correlates with the presence of the actin network. Indeed, tubes, as well as spikes, disappear where the actin network is destroyed ^6^ (Fig. 1, D and E and Materials and Methods). Moreover, the disappearance of tubes correlates with a change in membrane aspect, from rugged to smooth (Fig. 1D). A possible effect of membrane curvature induced by our SpVCA attachment is ruled out (Supplementary Information and Supplementary Fig. 2).

## Characterization of tubes

To assess where new actin monomers are incorporated during tube growth, we initiate actin assembly with Alexa568-labelled actin (red), and we incorporate new monomers of Alexa488-labelled actin (green) after 20 minutes (Materials and Methods). As previously observed for actin networks growing around polystyrene beads ^8,9^, new monomers insert at the liposome surface (Fig. 2A). Strikingly, new (green) monomers are also observed within the already grown (red) actin network (Fig. 2A), indicating new actin incorporation on the sides of membrane tubes (evidenced by phase contrast imaging, Fig. 2A, left), where SpVCA, the activator of actin polymerization, is also present (Fig. 2B).

**Figure 2:**
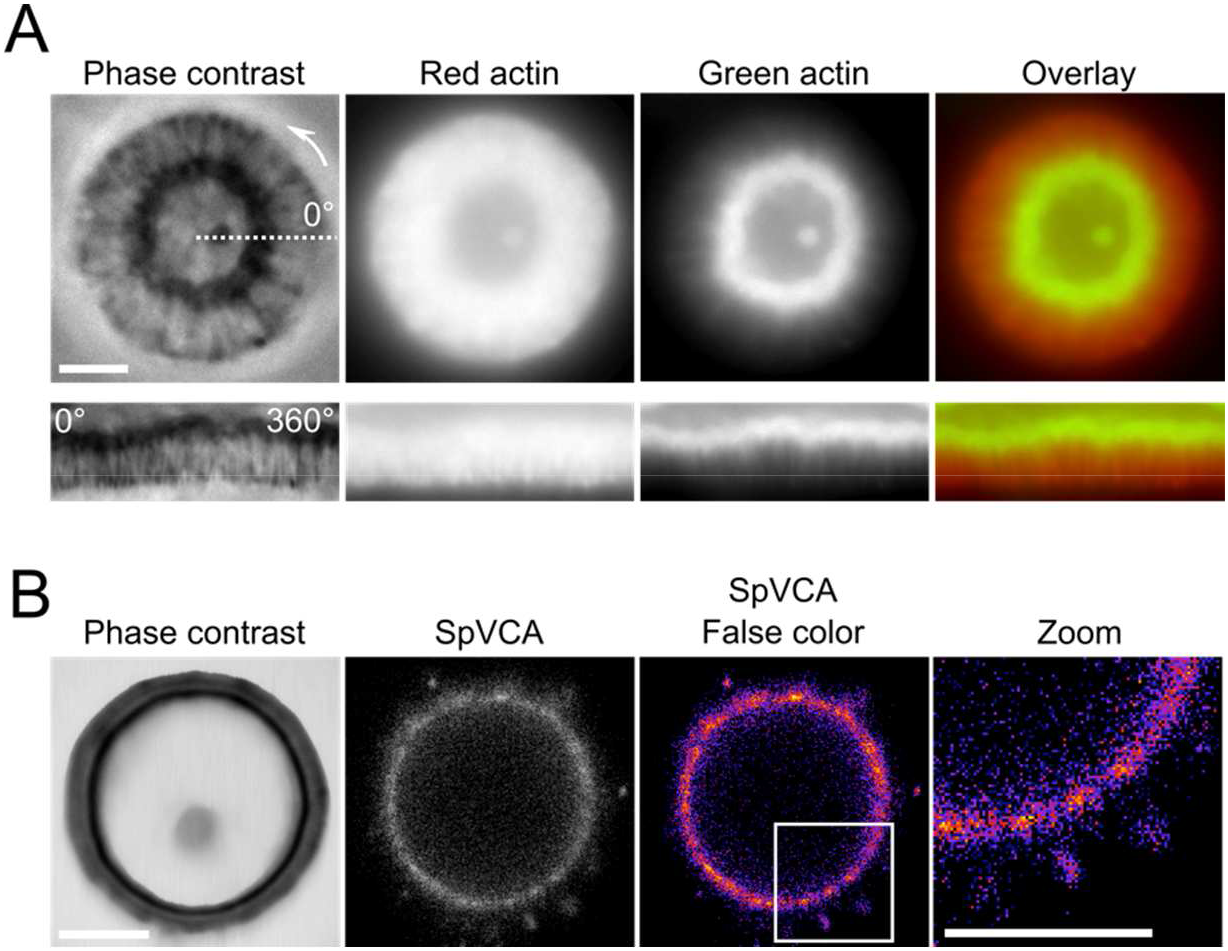
Actin incorporation during tube formation. (A) Top: a red actin network is grown for 20 minutes, then an excess of green actin is added, so green regions indicate newly polymerized actin. Bottom: corresponding polar plots. (B) Images of the activator of actin polymerization, SpVCA. False color image and zoom in (white rectangle). Phase contrast and epifluorescence microscopy of the actin network labeled with actin-Alexa568 (red) and actin-Alexa488 (green) in (A), and of SpVCA-Alexa546 in (B). Scale bars, 5μm.

We find that the average length of the longest tubes increases linearly with network thickness (Fig. 3, A and B). In fact, tube length roughly equals the thickness of the actin network, independent of the membrane tension (Fig. 3B, slope 0.89 ± 0.04). Moreover, we find that tubes grow simultaneously with the actin network (Fig. 3, C and D and Supplementary Fig. 3). An important observation is that there is a distribution of tube lengths within the actin network. Indeed, shorter tubes are present, since total fluorescence intensity decreases with distance from the liposome surface (Materials and Methods, Supplementary Fig. 4, A and B). Tubes shorter than the network thickness are clearly visible by confocal microscopy (Supplementary Fig. 4C).

**Figure 3:**
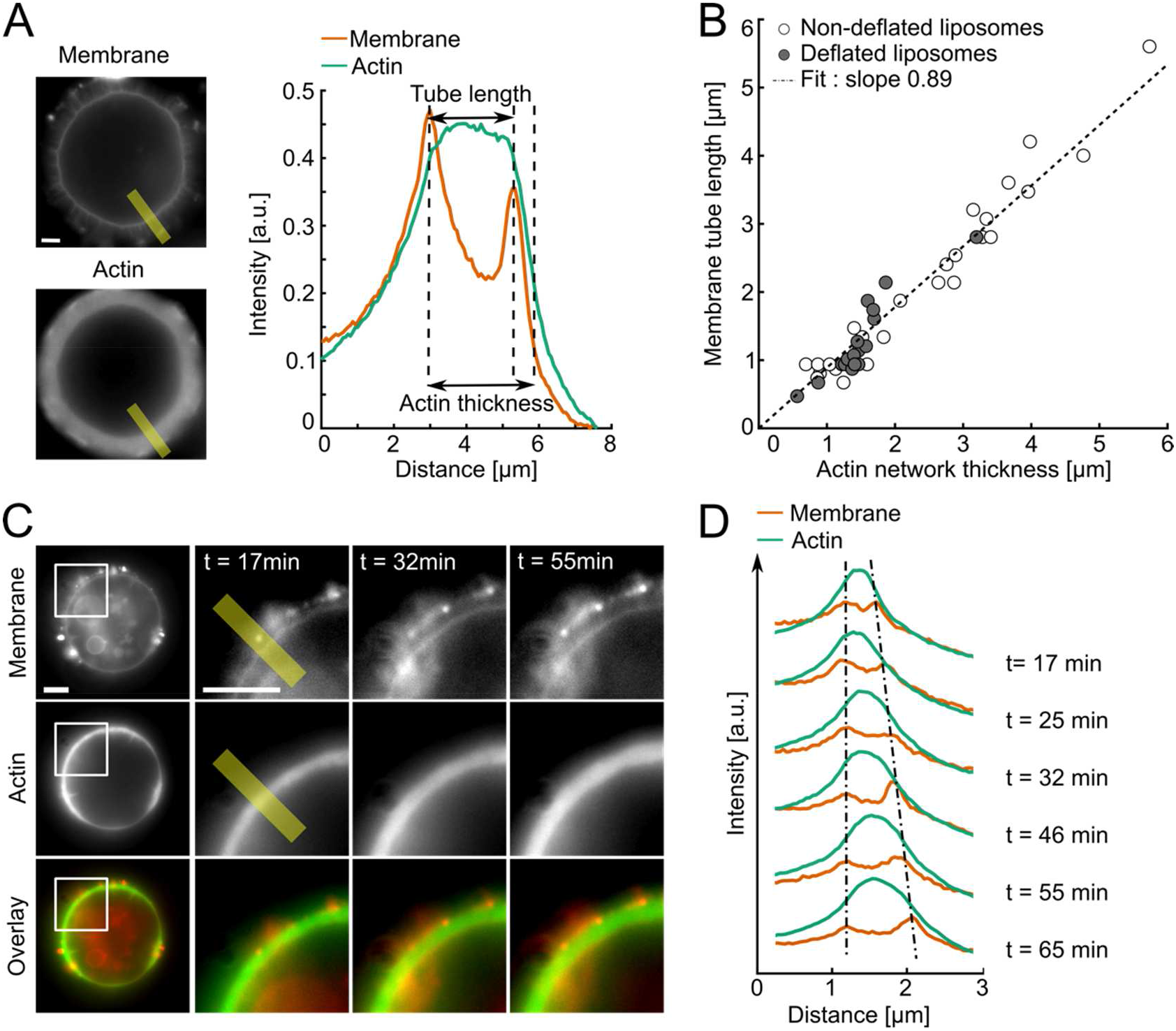
Tube length compared to network thickness. (A) Tube length and actin network thickness are measured from fluorescence intensity profiles (thick yellow line) of the membrane (red) and the actin (green) channels (Materials and Methods). (B) Tube length as a function of actin network thickness. White circles: non-deflated liposomes. Grey circles: deflated liposomes. (C) Dynamics of tube growth (times indicate elapsed time from the start of actin polymerization). (D) Fluorescence profile of the thick yellow lines shown in (C). Membrane and actin fluorescence intensities plotted over time (indicated). Other examples are shown in Supplementary Fig. 2. Epifluorescence microscopy of membrane and actin. Scale bars, 5μm.

The origin of the accumulation in membrane fluorescence detected at the tip of some of the longer tubes is unclear. We observe that SpVCA forms aggregates on membranes and sticks membranes together, even in the absence of actin (Supplementary Fig. 5). It is possible that small vesicles are attached via SpVCA to the membrane before polymerization starts and are pushed outward by actin growth. However, the presence of different tube lengths rules out that tubes could be only formed by pre-existing attached vesicles.

## Characterization of spikes

We find that new actin is incorporated at the tips of the spikes as well as at the sides (Fig. 4A), consistent with the localization of SpVCA (Fig. 4B). A clump of actin is observable at the base of the spikes (Fig. 4C). The thickness of the clump bears no clear correlation with the length of the spikes (Supplementary Fig. 6A), but slightly correlates with their width (Supplementary Fig. 6B). Spikes initially elongate with time until polymerization slows down, the basal width of spikes, however, remains roughly constant over time (Fig. 4D and Supplementary Fig. 6C).

**Figure 4:**
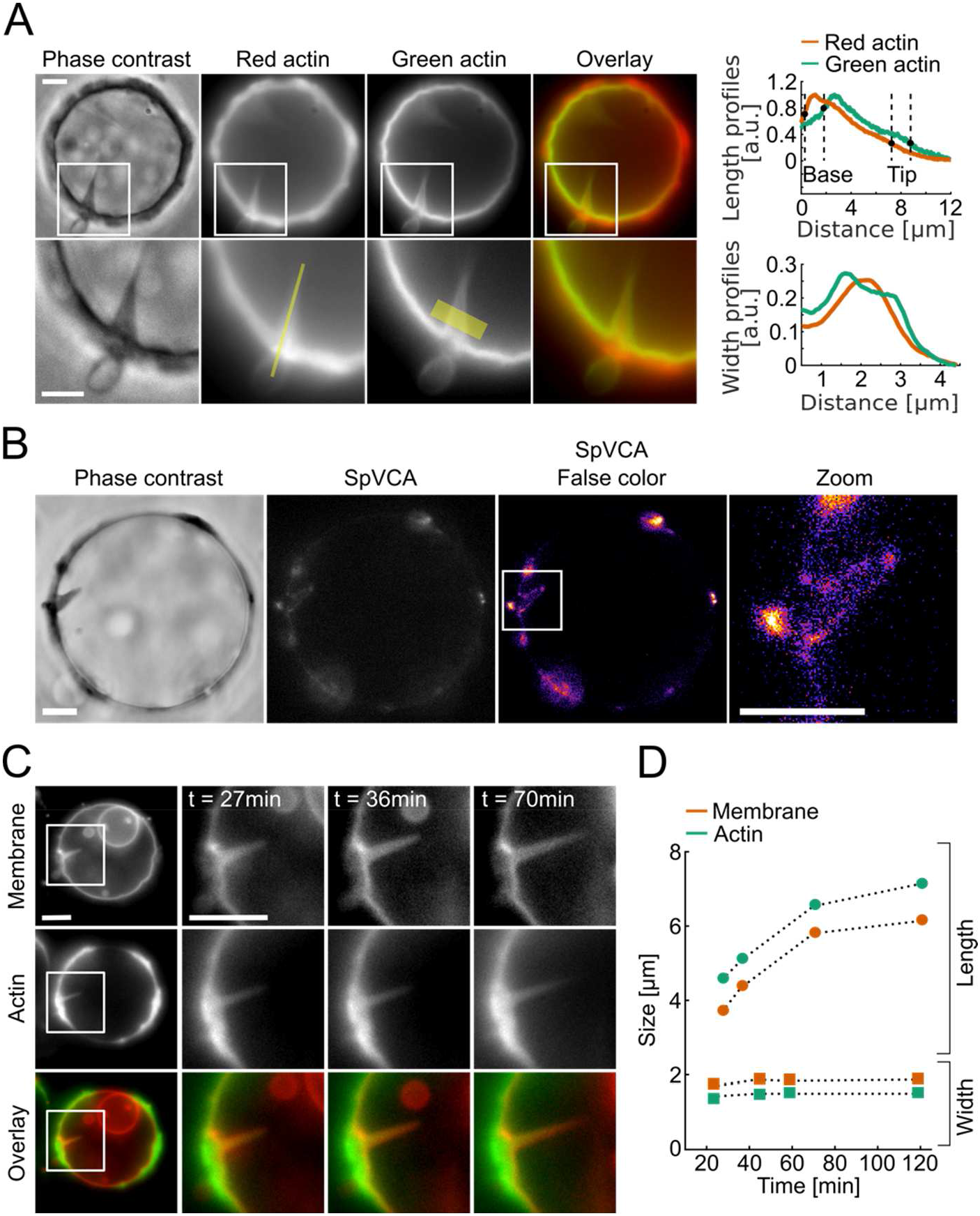
Actin incorporation in spikes. (A) Left: Two color experiment: green regions indicate newly polymerized actin. White squares, zooms. Right: fluorescence intensity profiles spike length (top, thin yellow line on zoomed image) and width (bottom, thick yellow line on zoomed image). (B) Activator of actin polymerization, SpVCA. White rectangle, zoom. (C) Dynamics of spike growth (time after actin polymerization starts). (D) Spike length and width over time, spike shown in C. Other examples in Supplementary Fig. 5. Dashed lines, guides to the eyes. Phase contrast and epifluorescence microscopy of the actin network (A), SpVCA-Alexa546 (B) membrane and actin network (C). Scale bars, 5 μm.

## Theoretical models for the growth of spikes and tubes

To rationalize the occurrence of spike-like structures arising solely from a uniformly polymerizing actin network, we use analytical modeling and numerical Finite Element calculations to evaluate the conditions under which large-scale membrane deformations may develop due to actin polymerization. We first estimate the normal stress exerted by the polymerization of an actin network (Material and Methods for details). The actin network can be modeled as a viscoelastic material with an elastic behavior at short time and a viscous behavior at long time due to network rearrangement, the cross-over time being on the order of 10 s ^10–12^. We choose to focus on the viscous behavior as the growth of the network occurs on timescales of tens of minutes.

We model the growth of the actin network by imposing a uniform actin polymerization velocity *v_p_* normal to the liposome membrane, a choice motivated by the dual color measurements in Fig. 4A. We solve the Stokes equation for the viscous gel polymerizing with a constant velocity normal to the liposome surface (Material and Methods). Actin polymerization on a flat membrane results in a uniform actin flow which does not generate any mechanical stress. A small perturbation of membrane shape modulates the actin velocity field which may generate viscous stress on the membrane. We show in the Material and Methods that the first order contribution to the normal stress exerted by the network on a periodically weakly deformed membrane, as illustrated in Fig. 5A, also vanishes identically. The lowest order contribution to the actin stress is quadratic with membrane deformation. This is in agreement with the finding that actin growing on a uniformly curved surface creates a normal stress proportional to the square of the curvature ^10,13^. In the case of a localized membrane perturbation, a Gaussian with amplitude *A* and width *b, u*(*x*) = *A* exp-(*x / b*)^2^ (Fig. 5B), we numerically calculate the normal stress exerted by an actin layer (Material and Methods). We obtain the pressure and velocity fields that arise in the actin layer (Fig. 5C). Velocity gradients in the growing actin layer, generated by the deformed surface, induce a normal force in the center of the perturbation, “pushing” the membrane inwards in the center of the perturbation (Fig. 5D). A scaling analysis of the Stokes equation, confirmed by our numerical calculation, shows that the normal stress *σ_nn_*, at the center of the perturbation (*x=0*) scales as *σ_nn_ ~ – ηA*^2^*b*^−3^*v_p_*, where *η* is the viscosity of the actin layer (Fig.5, E and F). It is important to realize that the normal stress exerted by the actin network on the membrane, when integrated over the area that surrounds the deformation, amounts to a zero net force. This contrasts with existing models of filopodia formation, which usually consider bundled actin filaments exerting a net pushing force on the membrane ^14^, invoking the friction force of treadmilling actin filament on the cellular actin cortex to balance this pushing force ^15^. While the latter approach might be appropriate to explain the physics of filopodia filled with bundled actin, here, we do not *a priori* distinguish the detailed structure of the actin network at the membrane from the one in the protrusion, treating the actin network as a continuum.

**Figure 5:**
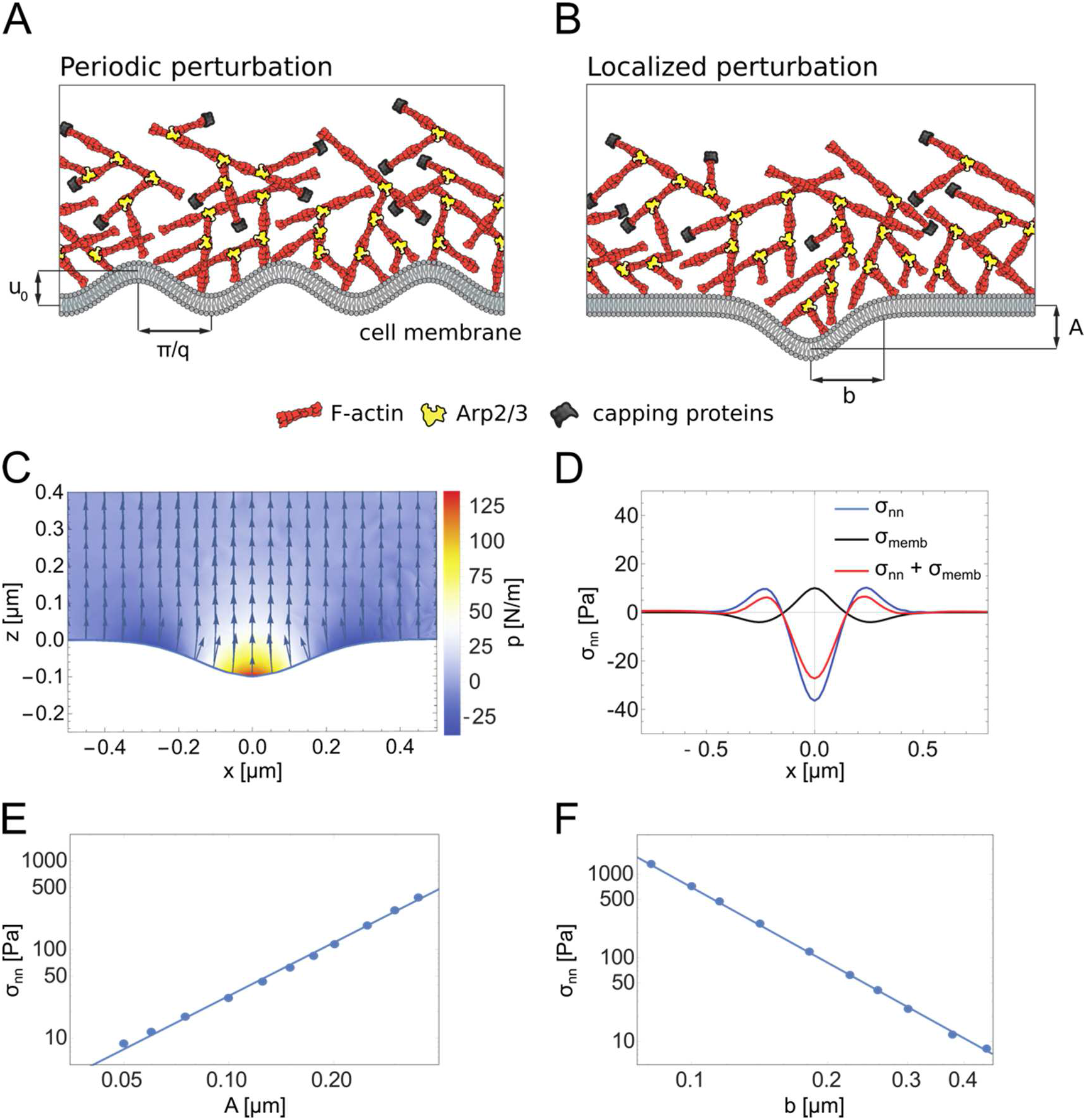
Model for spike initiation. Scheme of the initiation of a periodic (A) and localized (B) membrane deformation by the growth of the actin network. (C) Velocity field of a viscous network polymerizing over a membrane with a localized (gaussian) perturbation (amplitude *A=0.1 μm*, width *b=0.2μm*, polymerization velocity *v_p_=1 nm/s*, viscosity *η* = 10^4^ *Pa.s*). Color, pressure in the network layer. (D) Corresponding distribution of actin and membrane normal stresses (*σ_nn_* and *σ_memb_* respectively). (E, F) Scaling of *σ_nn_* as function of the amplitude for a value of *b= 0.22 μm* (E) and width for a value of *A=0.15μm* (F) of the perturbation (*v_p_= 1 nm/s*, viscosity *η*=10^4^ *Pa.s*).

The normal stress, on a deformable surface, is in our case balanced by the restoring elastic stress *σ_memb_* due to membrane elasticity. Neglecting the contribution of the membrane bending rigidity *κ* for simplicity, this stress corresponds to the membrane Laplace pressure *σ_memb_* = – *γC*, where *γ* is the membrane tension and *C~A/b*^2^ the local curvature (evaluated at the center of the localized perturbation). The balance of actin polymerization and membrane stresses defines a threshold amplitude *A*\* = *γb*/(*ηv_p_*). When the amplitude of the perturbation is smaller than this threshold (*A* < *A*\*) the membrane stress dominates and the perturbation relaxes. Above the threshold (*A* > *A*\*) the force exerted by the network is dominant and the instability develops. For *γ* ≈ 10^−6^*N/m* ^16^, *η* ≈ 10^4^*Pa s* (obtained through a scaling law from the elastic modulus *E* of the network and the viscoelastic time scale *τ_ve_* as *η* ≈ *Eτ_ve_*, where the elastic modulus *E* ≈ 10^3^*Pa* ^17^ and the viscoelastic timescale *τ_ve_* ≈ 10*s* ^11,12^) and *v_p_* ≈ 10^−9^*m/s*, the critical amplitude of a perturbation with a width *b* ≈ 10^−7^*m* is found to be *A*\* ≈ 10^−8^*m*. To evaluate whether such a perturbation could be reached by thermal fluctuations, the average membrane roughness corresponding to the fluctuation of a free membrane at thermal equilibrium is estimated as follows. With *ξ* the mesh size of the network, identifying (*A/b*)^2^ with the thermal average of the gradient of the membrane shape 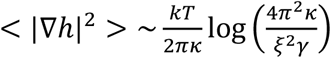 ^18^ and integrating over all wavelengths superior to the mesh size, we find a threshold tension for the appearance of spikes: 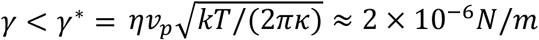. This value is in the range of membrane tension for liposomes under control conditions (*i.e*. non-deflated liposomes), but is larger than the tension of deflated liposomes, leading to the prediction that deflated liposomes are prone to the formation of spikes. In agreement with our experiments, the occurrence of spikes in nondeflated conditions (8.4%, Fig.1 C) is significantly lower than in deflated conditions (65.0%, Fig. 1C). A decrease of membrane tension has therefore a major influence on spike initiation.

We now develop a model for tube initiation. We consider a membrane deformation consisting of a very thin membrane tube connected to a flat membrane. The force required to pull a tube depends on membrane bending rigidity *κ* and membrane tension *γ* through 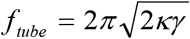, as extensively studied both theoretically and experimentally ^19,20^. Taking the bending energy of *10 kT* and a membrane tension of *γ*~10^−6^*N/m* ^16^, we find *f_tube_* ~2 pN. For the dynamic actin network to be able to pull a membrane tube, the tube force must be balanced by the mechanical stress in the growing network. Transient attachments between the membrane and the network exist when an actin filament is bound to the activator pVCA, as characterized experimentally ^21^. The growth of the actin network exerts a pulling force on this attachment site that can pull a membrane tube. Upon detachment, the tube retracts until one of the binding sites alongside the tube reaches the tube end, thus taking turn on the extraction force (Fig. 6A). This effectively results in the network exerting a long-lived localized force on the membrane, for times longer than the viscoelastic relaxation time, supporting a viscous description of the actin layer (Supplementary Information).

**Figure 6:**
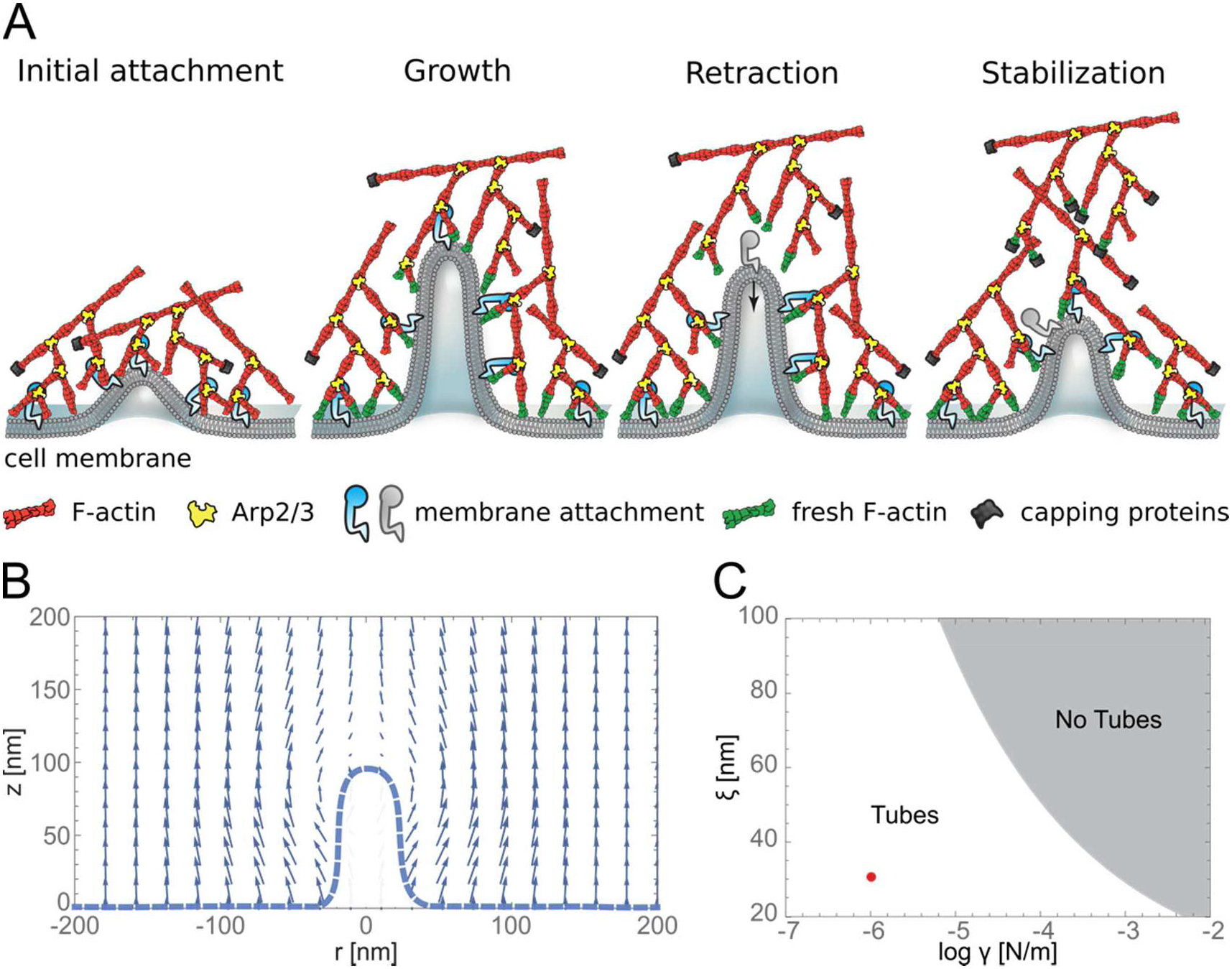
Model for tube initiation. (A) Scheme of a membrane tube pulled by the actin network and retraction due to detachment. (B) Velocity field of the actin network pulling the membrane tube. We assume a uniform polymerization *v_p_* at the liposome surface and model the presence of the tube as a disc with radius *r_tube_* = 20 *nm* and height *h* = 100 *nm*. (C) Phase diagram, mechanics of tube pulling as a function of mesh size *ξ* and membrane tension *γ*. Grey part, region where the viscous driving force is not sufficient to extract a tube (*f_tube_* = 2 *pN*, *κ* = 10*kT* and *h* = 140 *nm*). Red point, our experimental conditions.

The drag force exerted by the actin network (moving away from the liposome surface at a velocity *v_p_*) on the tip of the tube (moving at a velocity 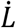) can be crudely estimated using the Stokes law: 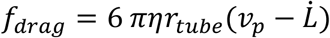 (Fig. 6B). At steady-state, the drag force balances the tube force, which provides the tube extraction velocity, 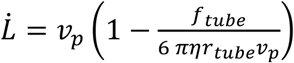. Tube extraction 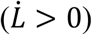 is therefore possible provided

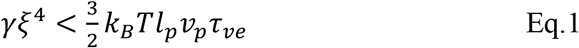

where we have used that *η* ≈ *Eτ_ve_* ≈ *k_b_Tl_p_τ_ve_/ξ*^4^ ^22^ (Fig. 6C). While a large enough membrane tension can in principle prevent tube extraction, the range of tension explored experimentally yields 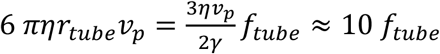 (with *v_p_* = 1*nm/s, η = Eτ_ve_* ≈ 10^4^Pa. s and *γ*~10^−6^N/m). Consequently, we find that membrane tubes can always be extracted by the growing actin network under the present conditions. Tube extraction however could in principle be prevented under high tension, or for a loose network (high value of the actin meshsize *ξ* > 100 *nm*). However, these regimes could not be explored due to experimental limitations of network growth under such conditions ^23^. Moreover the tube extraction velocity is very close to the actin polymerization velocity 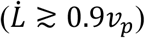. This explains why tubes initiated early during actin growth actually span the entire actin layer. The distribution in tube lengths inferred from Supplementary Fig. 4 can originate either from a distribution of tube nucleation time during the growth of the network or a distribution of rebinding time during tube retraction following a detachment from the actin network.

In yeast, actin is absolutely required for endocytosis, likely because of the high turgor pressure that opposes inward membrane deformations ^24–26^. The force needed to overcome the turgor pressure can reach 1000 pN ^27^, almost three orders of magnitude larger than the actin force in our *in-vitro* conditions. Using actin dynamics parameters relevant for yeast (polymerization velocity *v_p_* = 50*nm/s* ^1^ and actin network viscosity *η* = 2 10^5^*Pa. s* ^28^), the drag force generated by the actin network may indeed create the force required for membrane deformation leading to endocytosis (Supplementary Information).

## Discussion

The cell is a robust system where redundant mechanisms insure proper function, which makes detailed cell mechanisms difficult to decipher. This is true for membrane deformations into filopodia ^5^ or endocytic intermediates ^1^. Here, we show that a branched actin network growing at a membrane is able to mimic the initiation of either an endocytosis-like or a filopodia-like deformation. Endocytosis-like deformations appear to be a robust feature with regard to membrane tension whereas the initiation of filopodia-like structures is eased by a decreased membrane tension. Our results support recent findings that the initiation of dendritic filopodia and endocytosis primarily relies on the growth of a branched actin network ^1,3,5^.

Endocytosis is intimately dependent on the existence of a physical link between the actin network and the plasma membrane in yeast as well as in mammalian cells under high tension. Controlled endocytosis is abolished in yeast if this link is suppressed, although already endocytosed vesicles retain their extraordinary capacity to polymerize actin and even undergo actin-based motility ^3,29^. In our reconstituted system, in which actin nucleators are permanently linked to the liposome membrane, actin dynamics alone have the remarkable capacity to initiate endocytosis-like membrane deformations with a width smaller than, or of the order of, the actin mesh size.

A class of model for filopodia initiation assumes a particular actin organization in the protrusion, typically that of bundled actin filaments ^14,15,30,31^. Supported by our dual color actin measurements, our model for spike initiation assumes that actin polymerization occurs uniformly at the membrane, which indicates that new actin is incorporated all along the conical membrane surface, and not only at the tip of the protrusion as observed in Liu ^32^. Decreasing the membrane tension of the liposome decreases the critical amplitude for spike nucleation and increases the likelihood of spike formation (Fig. 5). This suggests a mechanism of spike formation different from that of tip growing protrusions, both in its initiation, and in its subsequent growth dynamics. Spikes are mimics of filopodia, especially in the case of dendritic filopodia whose formation relies on the Arp2/3 complex-branched network ^5^, as the suppression of the Arp2/3 complex system decreases the number of dendritic protrusions ^33^.

## Materials and Methods

### 1. Reagents, lipids, proteins

Chemicals are purchased from Sigma Aldrich (St. Louis, MO, USA) unless specified otherwise. L-alpha-phosphatidylcholine (EPC), 1,2-distearoyl-sn-glycero-3-phosphoethanolamine-N-[biotinyl polyethylene glycol 2000] (biotinylated lipids), 1,2-dioleoyl-sn-glycero-3-[[N(5-amino-1-carboxypentyl)iminodiacetic acid]succiny] nickel salt (DOGS-NTA-Ni) are purchased from Avanti polar lipids (Alabaster, USA). Texas Red^®^ 1,2-dipalmitoyl-sn-glycero-3-phosphocholine, triethylammonium salt is from Thermofisher. Actin is purchased from Cytoskeleton (Denver, USA) and used with no further purification. Fluorescent Alexa-488 actin and Alexa546 actin are obtained from Molecular Probes (Eugene, Oregon, USA). Porcine Arp2/3 complex is purchased from Cytoskeleton and used with no further purification. Biotin is purchased from Sigma-Aldrich (St. Louis, Missouri, USA), diluted in DMSO. Mouse α1β2 capping protein is purified as in ^34^. Untagged human profilin and SpVCA are purified as in ^7^. SpVCA is fluorescently labeled on the N-terminal amine with Alexa546 at pH 6.5 for 2 h at 4°C, desalted and then purified on a Superdex 200 column. His-pVCA-GST (GST-pVCA) is purified as for PRD-VCA-WAVE ^35^ and His-pVCA is essentially the same without the glutathione sepharose step. A solution of 30 μM monomeric actin containing 15% of labeled Alexa-488 actin is obtained by incubating the actin solution in G-Buffer (2 mM Tris, 0.2 mM CaCl2, 0.2 mM DTT, pH 8.0) overnight at 4°C. All proteins (SpVCA, profilin, CP, actin) are mixed in the isotonic or hypertonic working buffer. The isotonic working buffer contains 25 mM imidazol, 70 mM sucrose, 1 mM Tris, 50 mM KCl, 2 mM MgCl2, 0.1 mM DTT, 1.6 mM ATP, 0.02 mg/mL β-casein, adjusted to pH 7.4. The hypertonic working buffer differs only by its sucrose concentration and contains 25 mM imidazol, 320 mM sucrose, 1 mM Tris, 50 mM KCl, 2 mM MgCl2, 0.1 mM DTT, 1.6 mM ATP, 0.02 mg/mL β-casein, adjusted to pH 7.4. Osmolarities of the isotonic and hypertonic working buffers are respectively 200 and 400 mosmol, as measured with a Vapor Pressure Osmometer (VAPRO 5600). In case of experiments with DOGS-NTA-Ni lipids, all proteins are diluted in a working buffer containing 280 mM glucose, 10 mM HEPES, 0.5 mM DABCO, 100 mM KCl, 4 mM MgCl2, 1 mM DTT, 10 mM ATP and 0. 05 mg/mL β-casein.

### 2. Liposome preparation

Liposomes are prepared using the electroformation technique. Briefly, 10 μl of a mixture of EPC lipids, 0.1% biotinylated lipids or 5% DOGS-NTA-Ni lipids, and 0.1% TexasRed lipids with a concentration of 2.5 mg/ml in chloroform/methanol 5:3 (v/v) are spread onto indium tin oxide (ITO)-coated plates under vacuum for 2 h. A chamber is formed using the ITO plates (their conductive sides facing each other) filled with a sucrose buffer (0.2 M sucrose, 2 mM Tris adjusted at pH 7.4) and sealed with hematocrit paste (Vitrex Medical, Denmark). Liposomes are formed by applying an alternate current voltage (10 Hz, 1 V) for 2 h. Liposomes are then incubated with an activator of actin polymerization (SpVCA, 350 nM) via a streptavidin-biotin link for 15 min. Isotonic liposomes are used right away for polymerizing actin in the isotonic working buffer. To obtain deflated liposomes, an extra step is added: they are diluted twice in the hypertonic working buffer at 400 mOsmol and incubated for 30 min. The final solution is therefore at 300 mOsmol.

#### 2bis. Biotin-blocking experiments

SpVCA labeled with AlexaFluor546 and biotin are diluted in the isotonic working buffer and incubated for 10 min to reach final concentration of 350 nM SpVCA and various concentrations of biotin (87.5 nM, 175 nM, 262.5 nM, 350 nM). Note that 350 nM of biotin corresponds to a full saturation of the streptavidin sites of SpVCA. Unlabeled liposomes (99.9% EPC lipids, 0.1% biotinylated lipids) are then diluted twice in this solution and incubated for 15 min. Tubes and spikes are visualized by the fluorescence of SpVCA.

### 3. Actin cortices with a branched network

Condition 1: Actin polymerization is triggered by diluting the isotonic or deflated liposomes 6 times in a mix of respectively isotonic or hypertonic working buffer containing final concentrations of 3 μM monomeric actin (15% fluorescently labelled with AlexaFluor488), 3 μM profilin, 37 nM Arp2/3 complex, 25 nM CP. Note that the final concentrations of salt and ATP in both isotonic and hypertonic conditions are 0.3 mM NaCl, 41 mM KCl, 1.6 mM MgCl2, 0.02 mM CaCl2 and 1.5 mM ATP. Hypertonic conditions differ from isotonic conditions by 250 mM sucrose.

Condition 2: Same protocol as in Condition 1 with unlabeled monomeric actin, unlabeled liposomes (99.9% EPC lipids, 0.1% biotinylated lipids) and SpVCA labeled with AlexaFluor546.

In Figure 1, panel C, non-deflated liposomes n=311 are distributed as follows: 215 from 3 experiments in Condition 1 and 96 from 2 experiments in Condition 2. Deflated liposomes n=123 are distributed as follows: 92 from 2 experiments in Condition 1 and 31 from one experiment in Condition 2.

### 4. Photo-damage of the actin network

The actin network area to photo-damage is defined with a diaphragm. The area is illuminated for 15 s with a Hg lamp and a FITC filter cube and the illumination is repeated until actin is completely destroyed or at least no longer detectable by eye.

### 5. Two color experiment

Liposomes are first incubated with 350 nM SpVCA for 15 min. This solution is then diluted 3-fold into a mix of isotonic buffer containing 3 μM actin (15% Alexa568-labeled, red), 37 nM Arp2/3 complex and 25 nM CP. After 20 min of incubation in these conditions, the solution is diluted 3 times in a mix of same protein concentrations containing 15% Alexa488-labeled actin, green.

### 6. Cryo-electron microscopy

To prepare small liposomes, a mixture of EPC lipids and 0.1% biotinylated lipids with a concentration of 1 mg/mL in chloroform/methanol 5:3 (v/v) is dried and resuspended under vortexing in a buffer containing 25 mM imidazol, 1 mM Tris, 50 mM KCl, 2 mM MgCl2, 0.1 mM DTT, 1.6 mM ATP, 0.02 mg/mL β-casein. Liposomes are then incubated with SpVCA (350 nM) for 15 min and finally flash-frozen for cryo-electron microscopy. Images were recorded under low dose conditions with a Tecnai G2 Lab6 electron microscope operating at 200 kV with a TVIPS F416 4K camera and with a resolution of 0.21 Å/pixel.

### 7. Observation of liposomes

*Observation in 2D*: epifluorescence (GFP filter cube, excitation 470 nm, emission 525 nm; Texas red filter cube: excitation 545-580 nm, emission 610 nm-IR), phase contrast and bright-field microscopy are performed using an IX70 Olympus inverted microscope with a 100x or a 60x oil-immersion objective. Images are collected by a charge coupled device CCD camera (CoolSnap, Photometrics, Roper Scientific).

*Observation in 3D*: confocal and bright-field microscopy are performed using an inverted Confocal Spinning Disk Roper/Nikon with a 100x or a 60x oil-immersion objective and lasers with wavelengths of 491 nm for actin and 561 nm for lipids. A FITC filter cube (excitation filter: 478-495 nm/emission filter: 510-555 nm) and a TxRed filter cube (excitation filter: 560-580 nm/emission filter: 600-650 nm) are used to acquire respectively actin and lipids fluorescence. Images are collected by a charge coupled device CCD camera (CoolSnap HQ2, Photometrics, Roper Scientific).

*3D data*: Z-stacks are acquired using the software Metamorph on each wavelength with a z-interval of 0.5 μm.

### 8. Image analyses of liposomes, tubes and spikes

*Image analyses* are performed with ImageJ software and data are processed on Matlab. The thickness of the actin network and the length of tube membranes is obtained from fluorescence intensity profiles (Fig. 3A). The first peak of the membrane profile determines the liposome surface, the actin network thickness is the distance between this peak and the half width at half maximum of the actin fluorescence profile. The length of the membrane tubes is obtained as the peak-to peak distance of the membrane fluorescence profile. The size of spikes (length, width) and actin network is determined by the corresponding positions of the inflexion points. Fluorescence profiles in each case (membrane, actin) are fitted with a polynomial function. The first maximum and the second minimum of the fit derivative, corresponding to inflexion points of the profile, determine the membrane or actin edges. The size is then the distance between the two edges. From actin fluorescence profile, actin network thickness at the base of spike is defined as the distance between the first maximum and first minimum of the fit derivative.

*To determine whether shorter tubes are present* in addition to the easily visualized long ones, we measure the total fluorescence intensity of the membrane on an arc that is displaced along a radial axis *r* from close to the liposome surface to the external part of the network. We hypothesize that tubes maintain a constant diameter along their length, as is established for pure membrane tubes ^19^. In these conditions, if all tubes have the same length, the total intensity should show a plateau as a function of *r*, until falling off to zero at an *r* where there are no more tubes (Supplementary Fig. 4A). Conversely, the total intensity would decrease as a function of *r* if tubes of different lengths were present (Supplementary Fig. 4A).

### 9. Statistical analyses

All statistical analyses are performed using MedCalc software. N-1 Chi-squared test is used to determine the statistical significance. Differences among samples were considered statistically significant when p < 0.05.

### 10. Theoretical model for spike initiation

To calculate the stress exerted by a viscous network, polymerizing at a curved surface we consider an incompressible stokes flow, described by force balance and incompressibility, i.e., 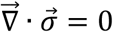 and 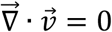, where 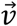 is the velocity of the gel and 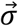 is the viscous stress in Cartesian coordinates, given by, 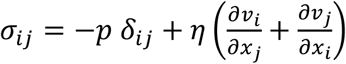. Polymerization of the actin network is encoded in this model by imposing the velocity of the network, normal to the surface of the curved interface. Moreover, we impose a stress free boundary condition at the outer layer, both for the normal as well as the tangential stress, i.e., *σ_nn_* = 0 and *σ_nt_* = 0. Note that, in the limit we consider, an infinite thick network, this corresponds to a uniform velocity in the z-direction.

We determine the first order correction of the normal stress on a deformed surface characterized by *u*(*x*) = *u*_0_ exp(*iqx*) along the *x* axis (*u*_0_ is the deformation amplitude and *q* the wave vector, Fig. 5A). We seek a solution for the velocity field within the network of the form *v_j_* = *v_j_*(*z*)exp(*iqx*), where the index *j* represents the coordinate *x* or *z*, and a pressure field of the form *p* = *p*(*z*)exp(*iqx*). Assuming that the network grows normal to the surface, the first order correction of the *x*-component of the velocity field satisfies the boundary condition *δv_x_*(*z* = 0) = –*v_p_∂_x_ u*(*x*) at the interface (*z=0*). We assume here a network of large thickness and require that the first order correction to the velocity vanishes at *z* → ∞. The first order corrections to the velocity and pressure in the network read *δv_x_*(*z*) = –*iqu*_0_(1 – *qz*)*v_p_*exp(–*qz*),*δv_z_*(*z*) = –*q*^2^*u*_0_*v_p_*z exp(–*qz*) and *δp*(*z*) = –2 *η q*^2^*u*_0_*v_p_*exp(–*qz*). At this order the actin normal stress turns out to vanish at any point of the liposome surface: *σ_nn_*(*x, z* = 0) = 2*η∂_z_v_z_* – *p* = 0. This implies that the membrane is linearly stable against small deformations in the presence of a growing actin network.

The second-order correction for the actin stress is in principle difficult to calculate, as the different modes of deformation are coupled. An analytical estimate can be obtained by expanding the surface normal vector up to second order, which yields the following scaling for the normal stress at the liposome surface, 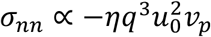. This weakly non-linear analysis reveals that there is a non-zero normal stress acting on the membrane, which we will later compare with the membrane contribution to address system stability. In order to get a numerical solution for the normal stress in a “localized” spike-like perturbation on the interface, as opposed to the periodic one presented above, we use a Finite Element Method from *Mathematics* with default settings. We implement a geometry as described in Fig. 5B, where the lower surface is parametrized with a Gaussian deformation as mentioned before, i.e, 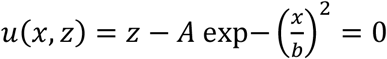 and we choose the height of the system to be much larger that the extend and amplitude of the perturbation (*h* = 2*μm*). Note that here, *b*, the characteristic lateral length of the localized perturbation, is related to the wavenumber *q*~1/*b* used for the linear analysis. To account for a constant polymerization, perpendicular to the lower surface we impose the velocity on the lower surface, i.e., 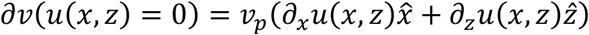, where *v_p_* is the normalized polymerization velocity and a vanishing normal and tangential stress at the upper boundary *z = h*, i. e., *σ_nn_*(*z = h*) = 0 and *σ_nt_*(*z = h*) = 0. Using this approach we could find the same scaling with amplitude and width of the perturbation, as found for the weakly non-linear analysis for a sinusoidal perturbation. Note also that here, by imposing the normal velocity at the interface, a choice that is motivated by the dual color images in Fig. 4A, we do not impose the tangential stress on the membrane, and hence this stress has to be balanced by an in-plane viscous stress in the membrane, which at this stage we do not model. These FEM simulations allow us to visualize the velocity field as well as the pressure throughout the network, indicating the increase in pressure inside the local perturbation caused by the local convergence of the velocity fields (Fig. 5C).

## Acknowledgments

We acknowledge Dr. Agnieszka Kawska at IlluScientia.com for the Fig.s. This work was supported by the French Agence Nationale pour la Recherche (ANR), grant ANR 09BLAN0283 and ANR 12BSV5001401, by Fondation pour la Recherche Médicale (FRM), grant DEQ20120323737, by the LabEx CelTisPhyBio postdoctoral fellowship (ML), No. ANR-10-LBX-0038 part of the IDEX PSL NANR-10-IDEX-0001-02 PSL, by Marie Curie Integration Grant PCIG12-GA-2012-334053, “Investissements d’Avenir” LabEx PALM (ANR-10-LABX-003 9-PALM), ANR grant ANR-15-CE13-0004-03 and ERC Starting Grant 677532. Our groups belong to the CNRS consortium CellTiss. This work was supported by grants from the French National Research Agency through the “Investments for the Future” (France-BioImaging, ANR-10-INSB-04), the PICT-IBiSA Institut Curie (Paris, France)

## Author contributions

CS, RK and VC have equal contributions. CS, VC performed experiments, analyzed data. RK performed the development of theoretical models. AA and MAG, JM, AdC, DL, CC, JP contributed to experimental data, ML and JFJ contributed to the development of the model, PS and CS designed the research. All authors contributed to write the paper.

## Author information

request for material should be addressed to PS and CS.

